# Chemometrics-based evaluation on effect of sonication, contact time and solvent-to-solid ratio on total phenolics and flavonoids, fatty acids and antibacterial potency of *Carica papaya* seed against *S. enteritidis, B. cereus, V. vulnificus* and *P. mirabilis*

**DOI:** 10.1101/2022.02.10.479904

**Authors:** Muhamad Shirwan Abdullah Sani

**Affiliations:** International Institute for Halal Research and Training, Ground Floor, Block EO, Kulliyyah of Engineering, International Islamic University Malaysia, PO Box 10, 50728 Kuala Lumpur, Malaysia; Konsortium Institut Halal IPT Malaysia, Ministry of Higher Education, Block E8, Complex E, Federal Government Administrative Centre, 62604 Putrajaya, Malaysia; The Catalytixs Solutions, No. 713, Jalan DPP ¼, Desa Permai Pedas, 71400 Pedas, Negeri Sembilan, Malaysia

**Keywords:** sonication-assisted extraction, contact time, solid-to-solvent ratio, antibacterial potency, *Carica papaya* seed

## Abstract

This study was aimed on extraction optimization of antibacterial agent from *Carica papaya* seed against *S. enteritidis, B. cereus, V. vulnificus* and *P. mirabilis* as affected by sonication-assisted extraction (SAE), contact time (CT) and solid-to-solvent ratio (SSR). The principal component analysis (PCA) and individual evaluation approaches identified that no SAE, 8 CT and 1:10 SSR was the best treatment with the highest antibacterial potency. The PCA identified no SAE, 8 CT, and 1:5 SSR as the second-beat treatment. The yield, total phenolic compound (TPC), C18:1n9t and C16:1 fatty acids (FAs) in no SAE, 8 CT and 1:10 SSR treatment inhibited the growth of *B. cereus, V. vulnificus* and *P. mirabilis* while C21:0 and C15:0 in 30 min SAE, 8 CT and 1:2 SSR inhibited the *S. enteritidis* growth. The yield, TPC, C18:1n9t and C16:1 FAs, and C6:0 and C24:1n9, C20:1, C4:0 and C20:0 FAs had antagonistic effects on *B. cereus, V. vulnificus* and *P. mirabilis* growths. The C21:0, C15:0, C6:0 and C13:0, and C23:0, C20:0 and C11:0 FAs had antagonistic effects on *S. enteritidis* growth. The PCA also denoted that the MIC_50_ and MIC_0_ had a higher variation than MIC; hence, the former variables were better to use in PCA.

## 1.0 Introduction

The antibacterial compounds had been searched from various plant by-products originating from tropical fruits such as pineapple, *Carica papaya*, and mango. *Carica papaya* cv. Sekaki or Hong Kong, Eksotika I and Eksotika II are varieties in Malaysia that provide an abundance of seed as a cheap source for the antibacterial product (Amazu et al., 2010). The commercial utilisation of the seeds is presently unknown, and data are still very scarce. They can also be sprinkled on salads or soups or added to any dish to substitute for black pepper (Malacrida et al., 2011). Reports on antibacterial activities from *Carica papaya* seeds are still scanty, as evident by the recent review on the agro-industrial potential of exotic fruit by-products (Sani et al., 2017b). The pioneering works by Emeruwa (1982) on epicarp (peel), endocarp and seeds of ripe *Carica papaya* against *Staphylococcus aureus, Bacillus cereus, Echerichia coli, Pseudomonas aeruginosa* and *Shigella flexneri*. Currently, Sani et al. (2021a) found that *S. enteritidis, B. cereus, V. vulnificus* and *P. mirabilis* were sensitive against a methanolic extract from the seed of riped *Carica papaya*. However, the optimisation information on extraction treatment to obtain the antibacterial potency from the *Carica papaya* seed extract is still negligible.

Generally, direct extraction without manipulating the extraction treatments is applied to study the antibacterial properties of natural by-products. Recently, new techniques such as microwave-assisted extraction (Mayer et al., 2008) and supercritical CO_2_ extraction (Zhao & Zhang, 2013) have been utilised to extract antibacterial compounds. However, relative to those techniques, the use of sonication-assisted extraction (SAE) is more convenient, affordable, environmentally friendly and industrially employed in local companies, by which the active compounds could be extracted in a shorter time and higher efficiency (Ya-Qin Ma et al., 2008). Time of contact (CT) is a necessary treatment to be optimised to minimise the process’s energy cost (Spigno et al., 2007). Longer contact time exposed active sites of the solid area, improved sample homogeneity (Chinn et al., 2011) and increased extraction yield (Romdhane & Gourdon, 2002). Solid-to-solvent ratio (SSR) was also reported to be a significant variation in the extraction of natural by-products (Pinelo et al., 2005) as it allowed maximum surface contact between solid and solvent (Zhang et al., 2007) and increased mass transfer rate (Pinelo et al., 2005). Both treatments also affected phenolic Wong, Tan, & Ho (2013) and flavonoid contents (Tan et al., 2011). Studies on contact time affecting antioxidative properties for fruit by-products such as citrus peel (Y.-Q. Ma et al., 2009) and SSR on grape seed (Pinelo et al., 2005) have been reported, but not for antibacterial properties.

A common approach to optimise the extraction of antibacterial compounds is by carrying out extraction treatment and evaluating the treatment effect individually, where only minimum inhibitory concentration (MIC) was considered. This approach brings minimal information for researchers to investigate the antibacterial activity of *Carica papaya* seed extract. Sani et al. (2017a) proposed including total phenolic compound and flavonoid compounds since these phytochemicals were reported to render antibacterial activities (Alonso-Esteban et al., 2019). However, to gain more information on the variables contributing to the antibacterial potency, fatty acids were included since 80.23% of *Carica papaya* seeds are dominated by the fatty acids and their esters (Sani et al., 2020). Therefore, our study quantified 37 fatty acids in their ester form and evaluated the effect of treatments and these variables using chemometric-based approachs i.e., principal component analysis (PCA) suitable for a multivariate dataset (Sani et al., 2021b). Very negligible report employed the chemometric-based approach to evaluate the antibacterial potency, especially on *Carica papaya* seed extract to date. It is also recommended to add MIC_50_ and MIC_0_ as additional antibacterial variables since these varaiables have variation that could render a more meaningful insight in the PCA results. It is anticipated that this study could provide a new insight about the antibacterial activities of *Carica papaya* seed extract, promote the application of cheap and ubiquitous by-products, and provide economic extraction procedures. This study will indirectly accelerate the quest for natural antibacterials.

## 2.0 Material and method

### 2.1 Experimental design

The experiment was designed as depicted in Figure 1. The experiment was divided into three effects which involved the effect of sonication-assisted extraction (SAE), contact time (CT) and solid-to-solvent ratio (SSR). The best treatment from each effect was applied in the following effect. For instance, the best treatment for the SAE effect was used in the CT effect, and both the best treatment from SAE and CT effect was then employed in the SSR effect. For each effect, the yield, total phenolic content (TPC) and total flavonoid content (TFC) of the extract ware determined as well as the antibacterial activities, consisting of minimum inhibitory concentration (MIC), MIC_50_, and MIC_0_ of *S. enteritidis*, *B. cereus, V. vulnificus* and *P. mirabilis*. Evaluation on the (1) individual treatment and variable and (2) treatments and multivariable via chemometrics were carried out and compared. The correlations among the variables were assessed, and the variables that characterised the treatment were identified.

**Figure 1:**
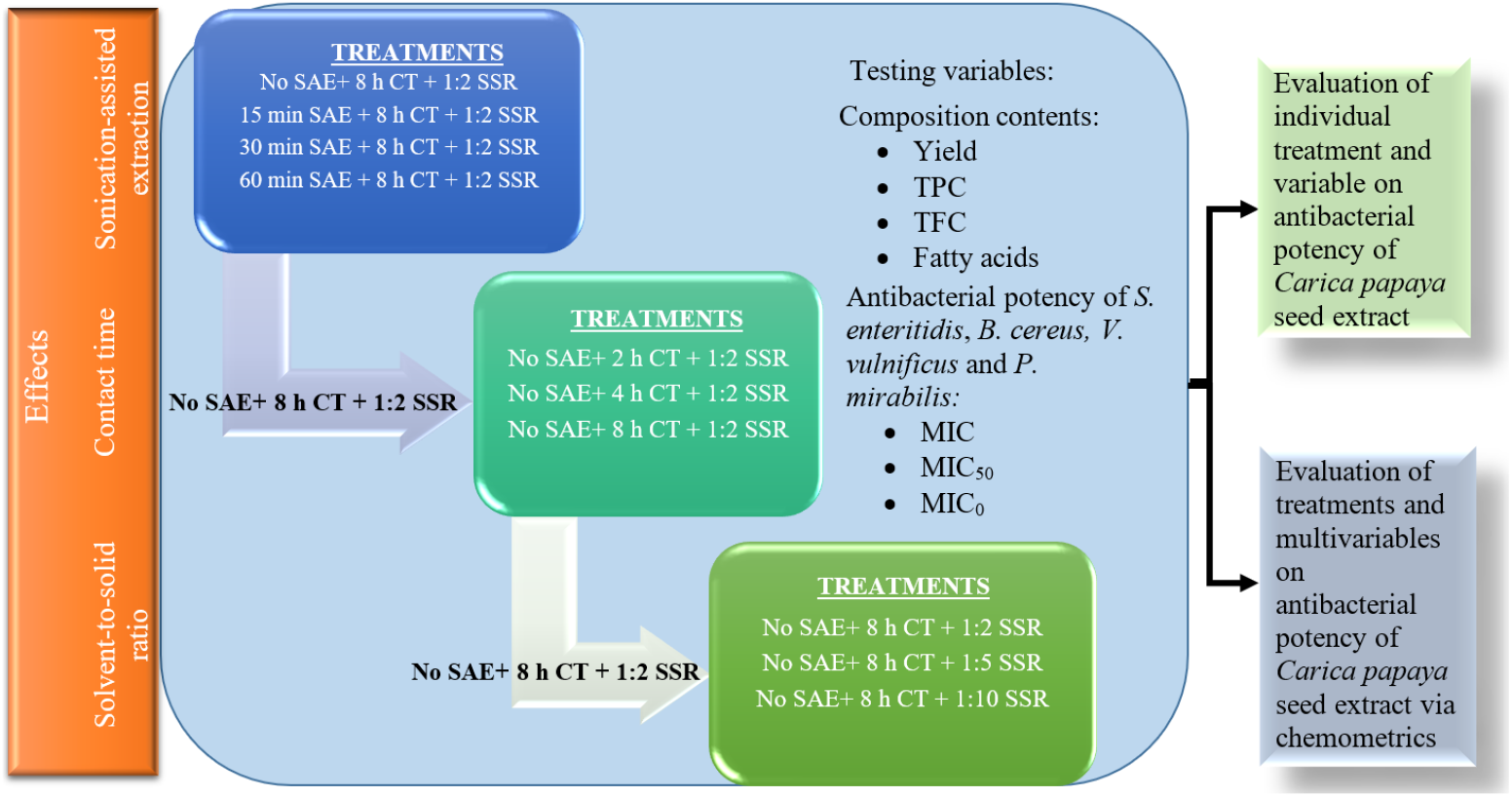
Experimental design of effect of sonication-assisted extraction, contact time and solvent-to-solid ratio on the antibacterial potency of *Carica papaya* seed against *S. enteritidis, B. cereus, V. vulnificus* and *P. mirabilis*

### 2.2 Plant material

*Carica papaya* cv. Sekaki fruit was bought from D’Lonek Sdn. Bhd. Organic Farm, Rembau, Negri Sembilan, Malaysia and numbered as SK 2368/14 voucher by Herbarium of Institute of Bioscience, Universiti Putra Malaysia. Carica papaya seeds were taken out from the fruit and treated as described by Sani, Bakar, Rahman, & Abas (2017).

### 2.3 Extraction of phytochemicals

Methanol (MeOH) was used as solvents in the extraction, according to Sani et al. (2020). No SAE (0 min SAE), 8 h CT and 1:2 SSR were standard extraction treatments. Briefly, 50 g of dried ground *Carica papaya* cv. Sekaki seed was weighed into a conical flask, and 100 mL of methanol was added. Extraction was carried out at 30°C in a shaker (100 rpm) followed by filtration through Whatman No.1 filter paper (GE Healthcare, UK). The filtrate was transferred into pre-weighed flat bottom flasks and concentrated using a rotary vacuum evaporator (Eyela N-1001, Japan) at 40°C.

To study the effect of SAE, extraction was carried out without SAE, and at 15 min, 30 min and 60 min sonication at 60 Hz with 550 W power. To study the effect of CT, extractions were carried out at 2, 4 and 8 h. To study the effect of SSR, extractions were carried out at 1:2, 1:5 and 1:10 SSR ratios. Each extraction treatment was done in triplicate.

### 2.4 Minimum inhibitory concentration (MIC), MIC_50_ and MIC_0_

A two-fold serial microdilution method of 96 multi-well microtiter plates was used for MIC determination with modifications. Briefly, 100 μL of tryptone soy broth (TSB) was added to each well. A volume of 100 μL of 0.5 × 10^5^ μg/mL crude extracts in dimethyl sulfoxide (DMSO) was added to the first well. A volume of 100 μl from the first test well was pipetted into the second well of each microtiter row, and then 100 μl of serial dilution was transferred from the second to the third and followed through until the eleventh well. A volume of 100 μL from the last well was discarded. An aliquote of 90 μL from each well was pipetted, mixed with 10 μL of 106 CFU/mL bacterial suspensions, and then added back into each well. This will make up 22.5 - 0.02 mg/mL extract concentrations from the first to the eleventh well. The microtiter plate was incubated at 37°C for 24 h on a Heidolph Inkubator and Titramax 1000 (Germany) at 210 rpm to prevent adherence and clumping, after which the optical density was measured at 600 nm in Tecan Infinite® 200 Microplate Reader (Switzerland) before (T_0_) and after (T_24_) incubation. A TSB medium incubated with a target bacterium (without an antibacterial agent) was used as a positive control of growth in the twelfth well in each row.

The MIC was defined as the lowest concentration of antibacterial agent showing a complete growth inhibition of the tested bacterial strains, which was related to a difference absorbance of T_24_ and T_0_ (T_24_-T_0_) equal to zero or negative values.

The graphs of percentage inhibition for each extraction treatment were plotted, and the MIC was compared to the percentage inhibition where MIC had 100% bacterial inhibition. MIC_50_ was determined by calculating the concentration that gave 50% inhibition by using linear regression (y = mx + c), where y = 50%, m = slope of regression, c = intercept of regression and x = concentration of extract at 50% inhibition. From the percentage inhibition graphs, the concentration of extract which gave 0% inhibition was also determined as MIC_0_. All determinations were performed in triplicate (Sowhini et al., 2020).

### 2.5 Quantitation of total phenolic content (TPC)

Total phenolic contents of *Carica papaya* crude extracts were determined by colourimetry assay with Folin-Ciocalteu according to Sani, Bakar, Rahman, & Abas (2017a). An amount of 0.05 g *Carica papaya* seed extract was diluted to 100 mL in a volumetric flask, and 1 mL of the diluted extract was mixed with 1 mL of 1:10 diluted Folin-Ciocalteu reagent (Sigma-Aldrich, Switzerland) in a 5 mL volumetric flask wrapped with aluminium foil and vortexed for 10 s. The mixture was then incubated at 30°C for 5 min, mixed with 1 mL sodium carbonate (10%, w/v) solution (Sigma-Aldrich, Switzerland) and marked up to the volume. Then, the mixture was vortexed (VTX-3000L, Copens Scientific, Germany) for another 10 s and incubated in the dark at 30°C for 30 min. The mixture produced a blue aqueous layer, and its absorbance was measured using a spectrophotometer (U-2810 Hitachi, Japan) at 747 nm against methanol as a blank in triplications. The exact incubation procedure was employed to prepare 0-10 mg/L gallic acid standard (Sigma-Aldrich, Switzerland) solutions in methanol. A calibration curve of gallic acid absorbance versus concentration was plotted, and the concentration of *Carica papaya* seed extract was computed from the calibration equation. Results were expressed as gallic acid equivalent (GAE) in mg/g of dry weight (DW) of the sample (mg GAE/g DW).

### 2.6 Quantitation of total flavonoid content (TFC)

A series of quercetin working standards at 0 - 12.5 mg/L, including ethanol as blank, was measured spectrophotometrically against ethanol at 438 nm, and a calibration curve was established. Total flavonoid contents of *Carica papaya* crude extracts were determined following Sani et al. (2017b). An amount of 0.05 g *Carica papaya* seed extract was diluted in a 100 mL volumetric flask with ethanol. A volume of 1.25 mL the diluted extract was mixed with 0.5 mL of 0.1 g/mL aluminium chloride solution (Sigma-Aldrich, Switzerland) and 0.5 mL of 1 M sodium acetate solution in 5 mL volumetric flask wrapped with aluminium foil, marked up to volume with ethanol and vortexed for 10 s. Then the solution mixture was incubated at 30°C for 15 min. After incubation, the absorbance of a yellow colour solution indicated the presence of the flavonoids was measured spectrophotometrically against ethanol. The measurements were carried out in triplicate, and results were expressed as quercetin equivalent (QE) per gram dry weight (DW) of the sample (mg QE/g DW).

### 2.7 Analysis of fatty acids methyl esters by gas chromatography-mass spectrometer

A series of working FAMEs standard in hexane ranging from 0.0005 – 3 mg/mL was prepared in 1 mL volumetric flask and injected into gas chromatography-mass spectrometry (GC/MS). A concentration of 0.01 g/mL *Carica papaya* seed extract was mixed with 0.6 mL of hexane and 0.4 mL of 1 M solution of sodium methoxide. Then, the mixture was vortexed for 30 s. A volume of 0.6 mL of top hexane layer was analysed by GC/MS for FAME quantification.

Separation and detection of FAMEs was carried out on an Agilent-Technologies 7890A gas chromatography (GC) system equipped with an Agilent-Technologies 5975 mass spectrometer (MS) system (Agilent Technologies, USA). The working standards and top hexane layer of the *Carica papaya* seed extracts were injected into an injector temperature maintained at 260°C. A volume of 1 μL of the standard and extracts was split at 1:10 ratio and eluted into the GC system by helium at 1 mL/min flow rate. The FAMEs were separated by an HP-88 capillary column (100 m x 0.25 mm, film thickness 0.20 μm) with an oven temperature program at (1) 150°C for 5 min, (2) heated to 240°C at 4°C/min and (3) held for 15 min. The separated FAMEs were eluted through MS transfer line and mass quadrupole set at 230°C and 150°C, respectively, ionised at 70 eV and detected by MS system at a mass range of m/z 20–700 units. The FAMEs detection and quantification were operated in scan and selected ion monitoring (SIM) modes.

A retention-time-lock mode of stable palmitic acid (C16:0) was executed to avoid changes in retention times in calibration curves due to column maintenance or column change. The FAMEs were identified by their retention times, comparison of their mass fragmentation patterns with standards from the National Institute of Standard (NIST) Mass Spectral 11 library and confirmation with the working standards. The linearity of the calibration curve was established and assessed, where the correlation coefficient R^2^ > 0.98 indicated an acceptable identification (Sani et al., 2021b).

### 2.8 Statistical analysis

Data were expressed as mean ± standard deviation of extraction yield, TPC and TFC and MIC_50_. One-way analysis of variance (ANOVA) with Tukey’s test was conducted using XLSTAT-Pro (2014) statistical software (Addinsoft, Paris, France) to determine the significant difference between the means at 95% confidence level (p < 0.05) for extraction yield, TPC and TFC and MIC_50_.

In this study, a principal component analysis (PCA) of Pearson correlation at α of 0.05 was employed to describe the correlation and distribution of significant chemical constituents and fatty acids towards the antibacterial potency of *Carica papaya* seed extract affected by the sonication, contact time and solvent-to-solid ratio. For antibacterial variables, the MIC, MIC_50_ and MIC_0_ for *S. enteritidis, B. cereus, V. vulnificus* and *P. mirabilis* were assigned as MICSE, MICBC, MICVV, MICPM, MIC_50_SE, MIC_50_BC, MIC_50_VV, MIC_50_PM, MIC_0_SE, MIC_0BC_, MIC_0_VV and MIC_0_PM, respectively. The dataset was transformed into independent variables known as principal components (PCs). Cumulative variability (CV) of two dimensional PCs entailing PC1 and PC2 were computed for Carica papaya seed extract profiling. The variables with strong, moderate and weak factor loading (FL) were identified. Based on these FLs, the variable correlations and their contributions to extraction treatments were assessed.

## 3.0 Results and Discussion

### 3.1 MIC, MIC_50_ and MIC_0_ as affected by sonication, contact time and the solid-to-solvent ratio of *Carica papaya* seed extract

The MIC test, a descriptive antibacterial method, had only given limited information on bacterial inhibition (Vigil et al., 2005) and inadequate comparison between extraction treatments. Thus, Patton et al. (2006) applied MIC, MIC_50_ (minimum concentration which gave 50% inhibition) estimation (Al-Habsi & Niranjan, 2012) and MIC_0_ (minimum concentration which gave 0% inhibition) information from the percentage inhibition to investigate the effect of extraction treatments on antibacterial properties of extracts.

The MIC, MIC_50_ and MIC_0_ of *Carica papaya* methanolic seed extract as affected by SAE against *S. enteritidis, B. cereus, V. vulnificus* and *P. mirabili*s growths were shown in Table 1. The Carica papaya seed concentration range used for all tested pathogens was 22.5 - 0.02 mg/mL. Among the tested pathogens, the *S. enteritidis, V. vulnificus and P. mirabilis* gave the lowest MIC (5.63 mg/mL). *B. cereus* had a MIC of 11.25 mg/mL as affected by 15, 30 and 60 min SAE and MIC of 5.63 mg/mL as affected by no SAE.

**Table 1:** MIC, MIC_50_ and MIC_0_ of *Carica papaya* methanolic seed extracts against *S. enteritidis, B. cereus, V. vulnificus* and *P. mirabilis*

| Treatment<sup>1</sup> | MIC (mg/mL) |  |  |  | MIC<sub>50</sub><sup>3</sup> (mg/mL) |  |  |  | MIC<sub>0</sub> (mg/mL) |  |  |  |
| --- | --- | --- | --- | --- | --- | --- | --- | --- | --- | --- | --- | --- |
|  | *S. enteritidis* | *B. cereus* | *V. vulnificus* | *P. mirabilis* | *S. enteritidis* | *B. cereus* | *V. vulnificus* | *P. mirabilis* | *S. enteritidis* | *B. cereus* | *V. vulnificus* | *P. mirabilis* |
| 15 min SAE | 5.63 | 11.25 | 5.63 | 5.63 | 3.66 ± 0.06<sup>c</sup> | 3.94 ± 0.05<sup>a</sup> | 3.78 ± 0.04<sup>c</sup> | 3.99 ± 0.06<sup>a</sup> | 1.41 | 1.41 | 0.02 | 0.70 |
| 30 min SAE | 5.63 | 11.25 | 5.63 | 5.63 | 3.80 ± 0.06<sup>b</sup> | 3.40 ± 0.11<sup>de</sup> | 3.61 ± 0.16<sup>c</sup> | 2.90 ± 0.20<sup>b</sup> | 1.41 | < 0.02 | < 0.02 | 0.02 |
| 1 h SAE | 5.63 | 11.25 | 5.63 | 5.63 | 4.05 ± 0.01<sup>a</sup> | 3.64 ± 0.13<sup>b</sup> | 3.74 ± 0.16<sup>c</sup> | 2.37 ± 0.34<sup>c</sup> | 1.41 | < 0.02 | 0.02 | < 0.02 |
| 2 h CT | 5.63 | 11.25 | 11.25 | 11.25 | 4.04 ± 0.03<sup>a</sup> | 3.85 ± 0.20<sup>a</sup> | 4.45 ± 0.24<sup>a</sup> | 2.48 ± 0.24<sup>bc</sup> | 1.41 | < 0.02 | 0.35 | < 0.02 |
| 4 h CT | 5.63 | 11.25 | 11.25 | 11.25 | 4.02 ± 0.08<sup>a</sup> | 3.56 ± 0.09<sup>bc</sup> | 4.01 ± 0.10<sup>b</sup> | 2.41 ± 0.31<sup>c</sup> | 1.41 | < 0.02 | < 0.02 | < 0.02 |
| 8 h CT<sup>2</sup> | 5.63 | 5.63 | 5.63 | 5.63 | 3.67 ± 0.01<sup>c</sup> | 3.43 ± 0.07<sup>cd</sup> | 3.65 ± 0.08<sup>c</sup> | 2.45 ± 0.37<sup>c</sup> | 0.70 | < 0.02 | < 0.02 | < 0.02 |
| 1:5 SSR | 5.63 | 5.63 | 5.63 | 5.63 | 3.07 ± 0.05<sup>d</sup> | 3.22 ± 0.06<sup>f</sup> | 3.35 ± 0.06<sup>d</sup> | 1.84 ± 0.45<sup>d</sup> | 0.35 | < 0.02 | < 0.02 | < 0.02 |
| 1:10 SSR | 5.63 | 5.63 | 5.63 | 5.63 | 2.87 ± 0.10<sup>e</sup> | 3.27 ± 0.10<sup>ef</sup> | 1.87 ± 0.25<sup>e</sup> | 2.37 ± 0.22<sup>c</sup> | < 0.02 | < 0.02 | < 0.02 | < 0.02 |

<sup>1</sup>15 min SAE - 15 min sonication assisted extraction; 30 min SAE - 30 min sonication assisted extraction; 1 h SAE - 1 h sonication assisted extraction; 2 h CT - 2 h contact time; 4 h CT - 4 h contact time; 8 h CT - 8 h contact time; 1:5 SSR - 1:5 solid-to-solvent ratio; 1:10 SSR - 1:10 solid-to-solvent ratio. <sup>2</sup>Extract was prepared without sonication at 1:2 solid-to-solvent ratio. <sup>3</sup>Means ± S.D. are from triplicate measurements.

The lowest MIC_50_ value for *S. enteritidis* and *P. mirabilis* was obtained from 15 min SAE and 60 min SAE, respectively, while 30 min SAE rendered the lowest MIC_50_ for *B. cereus* and *V. vulnificus* (Table 1). However, as we performed the significant test on these data, the MIC_50_ from these treatments were not significantly different from the no SAE treatment. The no SAE treatment gave the lowest MIC_0_ than SAE treatments for all tested pathogens (Table 1). Based on MIC, MIC_50_ and MIC_0_ comparisons among treatments, no SAE was the best treatment compared to other SAE treatments.

Table 1 exhibited the CT effect on percentage inhibition, MIC, MIC_50_ and MIC_0_ of *S. enteritidis, B. cereus, V. vulnificus* and *P. mirabili*s. The MIC values were higher at 2 h and 4 h contact time for *B. cereus, V. vulnificus* and *P. mirabilis* than 8 h CT, indicating that longer contact time produced a higher concentration of antibacterial compounds due to larger surface contact area between solvent and solute (Chinn et al., 2011). The MIC of these microorganisms also exhibited a strong correlation with contact time (Table 1). However, the 8 h extraction did not inhibit *S. enteritidis*.

The lowest MIC_50_ (Table 1) values for *S. enteritidis, B. cereus* and *V. vulnificus* were obtained from 8 h contact time, where significant differences of inhibitions were shown against *S. enteritidis* and *V. vulnificus* only. At 4 h contact time, *P. mirabilis* attained an insignificant difference of MIC_50_ as compared to 8 h contact time.

All contact time treatments provided < 0.02 mg/mL of MIC_0_ against *B. cereus* and *P. mirabilis*. *S. enteritidis* was the most sensitive against 2, 4 and 8 h contact time treatments where this microorganism showed MIC_0_ at 1.41 mg/mL for both 2 h and 4 h contact time, while at 8 h contact time, the MIC_0_ reduced to 0.70 mg/mL (Table 1). The 8 h CT also affected the *V. vulnificus* inhibition by reducing the MIC_0_ from 0.35 mg/mL to < 0.02 mg/mL. We conclude that the 8 h CT was the best CT treatment since it gave MIC, MIC_50_ and MIC_0_.

The MIC, MIC_50_ and MIC_0_ (Table 1) were tabulated from the percentage inhibition of respectively tested pathogens. All SSR treatments had not affected the MIC since all the MIC values of *S. enteritidis*, *B. cereus*, *V. vulnificus*, and *P. mirabilis* remained unchanged as per the SSR increment.

The insignificant difference of MIC_50_ of *B. cereus* was given by 1:10 SSR (3.27 ± 0.10 mg/mL) as compared to 1:5 SSR (3.22 ± 0.06 mg/mL) (Table 1). The 1:5 SSR had positive effect against *P. mirabilis*, by rendering significant difference (p < 0.05) of MIC_50_ (1.84 ± 0.45 mg/mL). For *S. enteritidis* and *V. vulnificus*, the significant differences (p < 0.05) of lowest MIC_50_ (2.87 ± 0.10 mg/mL) and (1.87 ± 0.25 mg/mL), respectively were obtained from 1:10 SSR, where both MIC_50_ of these microorganisms indicated strong correlation against SSR.

However, all SSR treatments did not influence *B. cereus*, *V. vulnificus* and *P. mirabilis*, where the MIC_0_ for these microorganisms remained at < 0.02 mg/mL. The MIC_0_ from 1:10 SSR also had presented the lowest value for all microorganisms (< 0.02 mg/mL). The 1:10 SSR had lowered the MIC_0_ of *S. enteritidis* to < 0.02 mg/mL from 0.70 mg/mL of 1:2 SSR (Table 1). The 1:10 SSR was chosen as the best SSR treatment compared to the 1:5 SSR and 1:2 SSR due to the lowest MIC_50_ and MIC_0_.

### 3.2 Yield, TPC and TFC as affected by sonication, contact time and solid-to-solvent ratio of *Carica papaya* seed extract

Extraction yields by different extraction treatments are shown in Table 2. The application of SAE resulted insignificant different (p < 0.05) in the extraction yields (19.12 - 21.57 mg/g) as compared to no SAE yield. This finding contradicted the yield of pomegranate seeds (Kalamara et al., 2014) and orange peel by Khan, Abert-Vian, Fabiano-Tixier, Dangles, & Chemat (2010). This occurrence was possibly due to the extraction reaching equilibrium before 15 min SAE contact time (Tian et al., 2013), reducing solvent’s permeability into cell structures on account of insoluble lipids existence on the ruptured cell (Tian et al., 2013) or re-adsorption of active components because of a large specific area of the ruptured cells (Dong et al., 2010). The extraction yield of *Carica papaya* seed extract also increased as contact time increased (Table 2) (Romdhane & Gourdon, 2002; Sargenti & Vichnewski, 2000; Spigno et al., 2007). Among the treatments, 8 h CT had produced the significant (p < 0.05) highest yield (21.59 mg/g). The yield of *Carica papaya* seed extract increased as SSR increased (Table 2), where 1:5 and 1:10 SSR showed significant yield (p < 0.05) as compared to 1:2 SSR. The 1:10 SSR treatment also indicated the highest (62.78 mg/g) and significant amount of TPC (p < 0.05). The high SSR increased the concentration gradient between the solid and the solvent (Zhang et al., 2007), enhanced diffusion rate, and allowed greater extraction of solids by solvent. Hence, no SAE, 8 CT and 1:10 SSR were the best treatments due to the highest yield. A calibration curve of TPC for *Carica papaya* seed extracts was established to obtain calibration equation y = 0.0827x + 0.0007 with coefficient determination (R2) of 0.9999. All SAE time Table 2 showed lower TPC (18.03 - 19.38 mg GAE/g DW) than no SAE (20.10 ± 0.74 mg GAE/g DW) and 60 min SAE. These treatments also exhibited insignificant difference (p < 0.05) of TPC as compared to no SAE, due to small energy generation during sonication, causing of low cavitation bubbles in the cell wall. This result was in agreement with Sargenti & Vichnewski (2000) work on Lychnophora ericoides and the recovery of phenolic compounds in honey (Biesaga & Pyrzyńska, 2013). Romdhane & Gourdon (2002) also achieved low TPC of woad seed (*Isatis tinetoria*) due to thermal dissociation of TPC during SAE (Luque-García & Luque De Castro, 2003; Y. Q. Ma et al., 2008).

**Table 2:** Extraction yield, total phenolic and total flavonoid contents of *Carica papaya* seed extracts

| Treatment<sup>1</sup> | Extraction yield<sup>3</sup> (mg/g) | TPC<sup>3</sup> (mg GAE / g DW) | TFC<sup>3</sup> (mg QE / g DW) |
| --- | --- | --- | --- |
| 15 min SAE | 20.73 ± 0.15<sup>b</sup> | 18.39 ± 0.81<sup>d</sup> | 2.31 ± 0.11<sup>d</sup> |
| 30 min SAE | 19.12 ± 0.15<sup>b</sup> | 18.03 ± 0.71<sup>d</sup> | 2.70 ± 0.03<sup>c</sup> |
| 1 h SAE | 21.57 ± 0.34<sup>b</sup> | 19.38 ± 0.74<sup>cd</sup> | 4.94 ± 0.18<sup>a</sup> |
| 2 h CT | 11.95 ± 0.07<sup>c</sup> | 9.85 ± 0.51<sup>f</sup> | 4.43 ± 0.31<sup>b</sup> |
| 4 h CT | 15.75 ± 0.08<sup>c</sup> | 13.89 ± 0.75<sup>e</sup> | 2.86 ± 0.15<sup>c</sup> |
| 8 h CT<sup>2</sup> | 21.59 ± 0.07<sup>b</sup> | 20.10 ± 0.74<sup>c</sup> | 1.91 ± 0.04<sup>e</sup> |
| 1:5 SSR | 76.02 ± 0.13<sup>a</sup> | 51.05 ± 1.99<sup>b</sup> | 2.32 ± 0.09<sup>d</sup> |
| 1:10 SSR | 81.15 ± 0.48<sup>a</sup> | 62.78 ± 1.51<sup>a</sup> | 2.20 ± 0.07<sup>d</sup> |

<sup>1</sup>15 min SAE - 15 min sonication assisted extraction; 30 min SAE - 30 min sonication assisted extraction; 1 h SAE - 1 h sonication assisted extraction; 2 h CT - 2 h contact time; 4 h CT - 4 h contact time; 8 h CT - 8 h contact time; 1:5 SSR - 1:5 solid-to-solvent ratio; 1:10 SSR - 1:10 solid-to-solvent ratio. <sup>2</sup>Extract was prepared without sonication at 1:2 solid-to-solvent ratio. <sup>3</sup>Means ± S.D. are from triplicate measurements.

The TPC of *Carica papaya* seed extract also increased as contact time increased (Table 2) (Romdhane & Gourdon, 2002; Sargenti & Vichnewski, 2000; Spigno et al., 2007). Among the treatments, 8 h CT had produced the significant (p < 0.05) highest yield (21.59 mg/g). The 8 h CT also exhibited the highest amount of TPC (20.10 mg/g) with a significant difference (p < 0.05) value as compared to other treatments due to longer contact time had improved surface area and slurry homogeneity of the sample; hence, provided a positive influence on phenolics (Chinn et al., 2011) by allowing the progressive release of phenolics from solid matrix to solvent (Spigno et al., 2007). Thus, the 8 h CT was the best CT treatment since it gave the highest values of yield and TPC and the lowest values of MIC, MIC_50_ and MIC_0_.

The calibration equation of TFC for *Carica papaya* seed extracts was established to obtain calibration equation y = 0.0762x - 0.0126 with an R^2^ of 0.9994. The SAE gave a significant difference (p < 0.05) of TFC (2.31 - 4.94 mg QE/g DW) in Table 2 as compared to no SAE treatment TFC (1.91 mg QE/g DW), where the 60 min SAE demonstrated the highest value, possibly due to flavonoids in the Carica papaya seed were in the form of flavonoid glycosides, which have thermal stability (Biesaga & Pyrzyńska, 2013; Ya Qin Ma et al., 2008). Since the *Carica papaya* seed used in this study did not undergo an acid hydrolysis process, only flavonoid glycosides were extracted out (Khoddami et al., 2013). Pan, Yu, Zhu, & Qiao (2012) claimed that the highest recovery of TFC in hawthorn seed was obtained at 91°C, which exceeded the temperature observed in this study (70°C), indicating the increment of temperature during SAE had not destabilised flavonoid glycosides. The increment of temperature during SAE was also reported to enhance solubility, increase the diffusion coefficient, and increase the extraction rate of TFC (Cacace & Mazza, 2003). From this finding, the no SAE was the best treatment compared to other SAE treatments.

On the other hand, the CT effect on TFC exhibited an inverse trend, where 2 h CT yielded the highest (4.43 mg/g) with a significant amount (p < 0.05). Even though Spigno et al. (2007) found that the optimum contact time for flavonoids in grape marc extraction was 5 h, our result showed the highest TPC, which was in line with the TPC of *Salvia officinalis* by Durling et al. (2007) at 8 h.

The 1:5 SSR produced the highest TFC (2.32 mg/g) with insignificant difference (p < 0.05) compared to the 1:10 SSR (2.20 mg/g). Although the TFC generally increased significantly (p < 0.05) for 1:10 and 1:5 SSR than the 1:2 SSR in Table 2, the TFC increment may not be directly proportional (Tan et al., 2011) since the TFC increment was halted once reaching the solid and solvent equilibrium (Pinelo et al., 2005) regardless of the SSR (Wong et al., 2013). Hence, the 1:10 SSR was the best SSR treatment compared to the 1:5 SSR and 1:2 SSR due to the TFC.

### 3.3 Correlation of total phenolic and flavonoids, fatty acids and antibacterial potency of *Carica papaya* seed extract on the extraction treatments

The purposes of using PCA in this study were to describe the correlation and distribution of yield, TPC, TFC and fatty acids on the antibacterial potency of *Carica papaya* seed extracts as affected by the SAE, CT and SSR.

Table 3 shows the characteristics of the analytical curves with the R^2^ values. The R^2^ > 0.98 indicated that the analytical curve values had established linear regression models, which were adequate for the FAMEs determination in the *Carica papaya* seed extract (Sani et al., 2021b). By analysing the FAMEs that take about 80.23% of the Carica papaya seed extract (Sani et al., 2020), we could investigate their influences on the yield, TPC and TFC on the pathogens inhibition via a chemometric technique such as PCA.

**Table 3:**
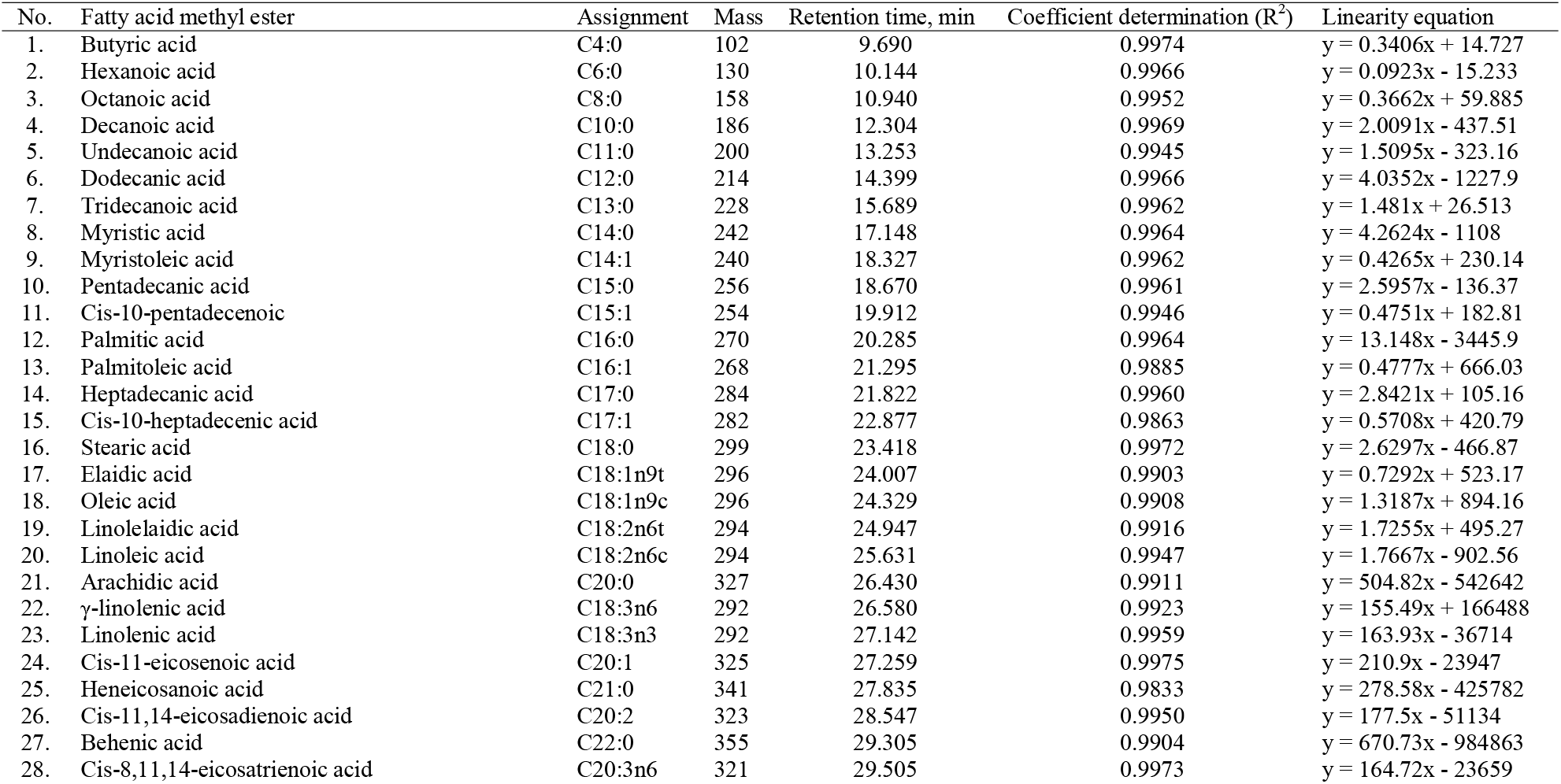
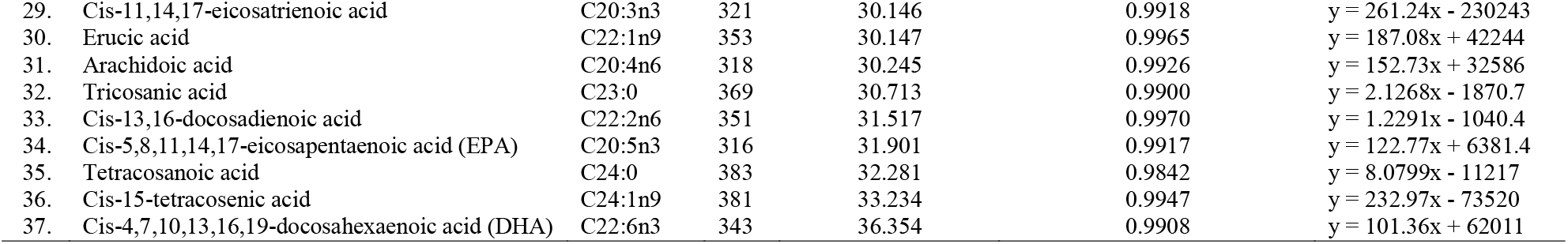
Characteristics of analytical curves for fatty acid methyl esters

The PCA exhibited two principal components (PCs) entailing 36 variables that represent cumulative variability of 46% with an eigenvalue (EV) of 6.57 in Figure 2 (a) for the whole dataset. In principle, variables far away from the axes F1 and F2 had a strong factor loading (FL). Of the 36 variables, the yield, TPC, C18:1n9t, C15:0, C16:0, C18:2n6c and C21:0, MIC_50_SE, MIC_50_VV, MIC_0_SE, had strong FL (FL ≥ |0.75|) that were dominant in this study. Besides, the TFC, C14:0, C16:0, C16:1, C18:0, C18:1n9c, C24:1n9, C6:0, C14:0, C18:0, and C23:0, MICBC, MICVV, MICPM, MIC_50_BC, MICBC, MIC_0SE_ had moderate FL (|0.500| < FL < |0.749|) while the other variables had a weak FL (FL ≤ |0.499|).

**Figure 2:**
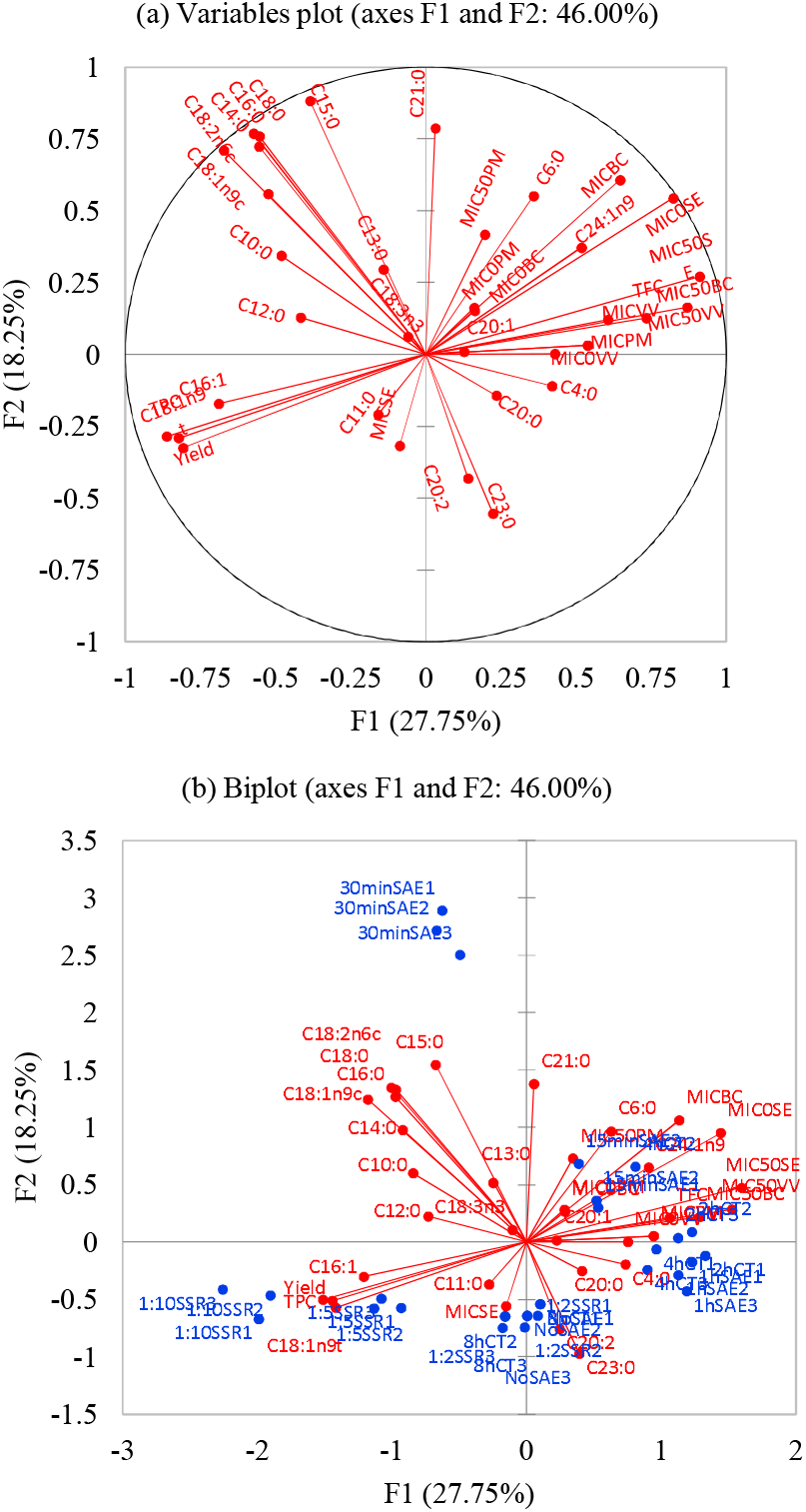
(a) Variable plot and (b) biplot of correlaton among the yield, TPC, TFC and fatty acids and antibacterial potency of *Carica papaya* seed extract

Figure 2 (a) of the variable plot also depicts the positive correlations among yield, TPC, C18:1n9t and C16:1, since these variables were positioned together. Although fatty acids and FAMEs were dominant in *Carica papaya* seed extract, other compounds such as organic acids, fatty aldehydes, sterols, nitriles and amides that had a phenolic backbone may render the antibacterial potency (Sani et al., 2020); therefore, denoted the high TPC in this study. The yield, TPC, C18:1n9t and C16:1 also had a negative correlation with the antibacterial variables except for MICSE on the opposite side of Figure 1 (a), i.e., MIC_50_SE, MIC_50_VV, MIC_0_SE, MIC_50_PM, MIC_0_PM, MIC_0_BC, MICBC, MIC_50_BC, MICVV, MICPM and MIC_0_VV. This correlation signified that higher yield, TPC, C18:1n9t and C16:1 possibly lead to lower MIC, MIC_50_ and MIC_0_; hence, inhibited the growth of *B. cereus, V. vulnificus* and *P. mirabilis*. Likewise, since the C6:0 and C24:1n9 were located in the same quadrant of these antibacterial variables, they were moderately rendered increment growth of the pathogens. Also, the C20:1, C4:0 and C20:0 had weak effects on the increment of pathogens’ growth. The opposite direction of the yield, TPC, C18:1n9t and C16:1 against C6:0 and C24:1n9, C20:1, C4:0 and C20:0 also indicated that these variables had antagonistic effects on the potency of *Carica papaya* seed extract. Hence, partially purified *Carica papaya* seed extract or a mixture of phenolics and purified C18:1n9t and C16:1 shall be employed instead of the crude extract to enhance the antibacterial potency of the *Carica papaya* seed extract against *B.cereus*, *V.vulnificus*, and *P. mirabilis*. This was evident since organic acids and sterols from *Carica papaya* seed extract have rendered MIC at 2.81 mg/mL, 0.35 mg/mL, and 1.41 mg/mL towards *B. cereus, V.vulnificus*, and *P. mirabilis*, respectively (Sani et al., 2021a). An interesting observation showed that C15:0, C18:0, C16:0, C14:0, C18:2n6c, C18:1n9c, C23:0 and C20:2 had no effect on the inhibition of B.cereus, V.vulnificus and P. mirabilis. This finding was due to their 90° direction towards the MIC, MIC_50_ and MIC_0_ of these pathogens.

Further investigation on the antibacterial variables, only MIC_50_SE, MIC_50_VV and MIC_0_SE had the strong FL (Figure 2 (a)) due to PCA measures variances in the dataset. This variance measurement fit for continuous variables compared to the end-point MIC value of pathogens (Sowhini et al., 2020). Hence, recording the antibacterial activity of *Carica papaya* seed extract would be better in MIC_50_ and MIC_0_ instead of MIC.

To note that C21:0 and C15:0 had strongly inhibited the *S. enteritidis* growth, while C6:0 and C13:0 had moderate effects in Figure 2 (a). On the contrary, C23:0 had moderately enhanced the growth of this pathogen, while C20:0 and C11:0 had weak influences. Also, the C10:0, C12:0, C4:0 and C20:0 had no inhibition effect on this pathogen. These variables also denoted the synergistic effect among the fatty acids and antagonistic reactions between C21:0, C15:0, C6:0 and C13:0, and C23:0, C20:0 and C11:0. This finding may propose that although individual short, medium, and long-chain fatty acids have antibacterial activity (Bae & Lee, 2017), their combination may also facilitate pathogen inhibition.

The biplot in Figure 2 (b) exhibited four specific clusters, i.e., 1:10 SSR, 1:5 SSR, 30 min SAE and 15 min SAE, and two mixed clusters. The 1:10 SSR was positioned at the most left of the F1 axes, followed by the 1:5 SSR where the yield, TPC, C18:1n9t and C16:1 were high while C6:0, C24:1n9 and C20:1 were low in these clusters. The extraction of *Carica papaya* seed through no SAE, 8 h CT and 1:10 SSR would be the best treatments to exert the highest antibacterial potency because of their opposite position of the antibacterial variables except MICSE. Likewise, the no SAE, 8 h CT and 1:5 SSR extraction treatments could be the second-best option.

The 30 min SAE cluster was also dominant as it is located to the left side of the F2 axes in Figure 2 (b). High C21:0 and C15:0 and low C20:2 and C23:0 characterised this cluster which also denoted that the extraction treatments 30 min SAE, 8 h CT and 1:2 SSR could inhibit the *S. enteritidis* as depicted with the opposite position of this cluster against the MICSE.

Nevertheless, the 15 min SAE cluster was positioned with the antibacterial variables except for MICSE, indicating 15 min SAE, 8 CT, and 1:2 SSR could not enhance the antibacterial potency of *Carica papaya* seed extract (Figure 2 (b)). This was due to high C6:0, C24:1n9 and C20:1 and low yield, TPC, C18:1n9t, and C16:1 may render antagonistic effects and had facilitated the pathogens’ growth. One mixed cluster consisting of 1h SAE, 2 h CT and 4 h CT also had indicated they could not inhibit the pathogens’ growth. In the same position, the C4:0 and C20:0 were dominant in this cluster. Another mixed cluster had the extracts produced from the standard extraction treatments, i.e., no SAE, 8 h CT and 1:2 SSR. The C23:0, C20:2, and C11:0 were high in this cluster, but they could not inhibit the *S. enteritidis*. To note, the two mixed clusters and 15 min SAE cluster were located at the centre of the F1 and F2 axes; therefore, they were the least significant extraction treatment in this study.

From these evaluations, the no SAE, 8 h CT and 1:10 SSR would be the best treatments to inhibit the *B.cereus, V.vulnificus*, and *P. mirabilis* growths while 30 min SAE, 8 h CT and 1:2 SSR could be the best extraction treatments to inhibit the *S. enteritidis*. It is also recommended to apply PCA to facilitate the interpretation of multivariable dataset instead of evaluating individual variables (Sani et al., 2021b). Besides providing information on the best extraction treatments, the PCA presents in-depth information on the variable correlations and proposes variables with synergistic and antagonistic effects that the evaluation of individual variables could not provide.

## 4.0 Conclusion

The chemometrics-based evaluation via PCA was superior over the individual evaluation treatment on antibacterial potency of *Carica papaya* seed extract. The no SAE, 8 h CT and 1:10 SSR were the most efficient treatments for extracting antibacterial compounds from Carica papaya seed than other SAE and CT treatments due to the highest yield and TPC. These treatments also gave the lowest MIC, MIC_50_ and MIC_0_ of *S. enteritidis, B. cereus, V. vulnificus* and *P. mirabilis*. Only *B. cereus* and *P. mirabilis* were sensitive against no SAE, 8 h CT and 1:10 SSR. All tested pathogens were sensitive against 1:10 SSR. Besides identifying the best treatment, the PCA had successfully identified the characterising variables on the treatments and synergistic and antagonistic effects of the variables. Although the PCA could delineate this information, only 46% of the dataset was explained in this study; hence, more variables and samples will be included in future research. The results and methods described here could help optimise extraction procedures through antibacterial test assessment and thus help reveal novel antibacterial compounds, which are expected to serve as an antibacterial agent food preservative.

## 5.0 Acknowledgement

This work was supported by the Malaysia Fundamental Research Grant Scheme (FRGS/1/2018/STG04/UIAM/03/1) of Ministry of Higher Education Malaysia.

## 6.0 Conflict of interest statement

We declare no conflict of interest.

## 7.0 Research involving human participants and/or animals

We declare no human and/or animals involved in this study.

## Notes

### Competing Interest Statement

The authors have declared no competing interest.

